# Optimizing Multi-Environment Trials in The US Rice Belt via Smart-Climate-Soil Prediction Based-Models and Economic Importance

**DOI:** 10.1101/2024.07.02.601777

**Authors:** Melina Prado, Adam Famoso, Kurt Guidry, Roberto Fritsche-Neto

## Abstract

Rice breeding programs globally have worked to release increasingly productive and climate-smart cultivars, but the genetic gains have been limited for some reasons. One is the capacity for field phenotyping, which presents elevated costs and an unclear approach to defining the number and allocation of multi-environmental trials (MET). To address this challenge, we used soil information and ten years of historical weather data from the USA rice belt, which was translated into rice response based on the rice cardinal temperatures and crop stages. Next, we eliminated those highly correlated Environmental Covariates (ECs) (>0.95) and applied a supervised algorithm for feature selection using two years of data (2021-22) and 25 genotypes evaluated for grain yield in 18 representative locations in the Southern USA. To test the trials’ optimization, we performed the joint analysis using prediction-based models in four different scenarios: I) considering trials as non-related, ii) including the environmental relationship matrix calculated from ECs, iii) within clusters; iv) sampling one location per cluster. Finally, we weigh the trial’s allocation considering the counties’ economic importance and the environmental group to which they belong. Our findings show that eight ECs explained 58% of grain yield variation across sites and 53% of the observed GxE. Moreover, it is possible to reduce 28% the number of locations without significant loss in accuracy. Furthermore, the US Rice belt comprises four clusters, with economic importance varying from 13 to 45%. These results will help us better allocate trials in advance and reduce costs without penalizing accuracy.

## 1. Introduction

Among the core objectives of rice breeding programs is the release of cultivars with improved yield, nutritional capacity, resistance to pests/diseases, and climate-smart (Hickey et al., 2019). However, some aspects hold back the genetic gains of breeding programs. One is the high costs associated with phenotyping (Araus & Cairns, 2014; Furbank & Tester, 2011), which can cause uncertainty in the number and allocation of trials throughout the locations to increase further. Another is unclear approaches to align multi-environmental trials (MET) with the target population of environments (TPE), especially in a climate change scenario that constantly challenges the definition of a TPE (Cooper & Messina, 2021, 2021).

Design of TPEs generally considers historical data from the soil, climate, hydrological aspects, management strategies, and sometimes socioeconomic data from locations where the crops are frequently produced, that is, it represents a set of characteristics that constitute future growing seasons, years and environments (Cooper et al., 2023; Crespo-Herrera et al., 2021). To avoid the yield gaps between the expected yield potential in the TPE and the real on-farm yield that farmers achieve, it is important to characterize better and select the locations used for testing/selection and to understand how they are related (mega-environments) by a process called enviromic or environtyping. Specifically, in this process, the environmental covariates are collected and processed in appropriate MET groupings, which are analyzed concerning the alignment with TPE and, thus, used to capture genotype reaction norms models (Callister et al., 2024; Cooper & Messina, 2021; Costa-Neto et al., 2023).

The concepts of environmental characterization, the genotype by environment interaction (GxE), and the target population of environments have been mentioned for many years in corn breeding in the USA, with this information presenting itself as a valuable resource for the decision-making process in breeding programs (Boer et al., 2007; Gaffney et al., 2015). However, little attention has been given to the American rice belt’s environmental characterization and TPE delineation, which differ in environmental effects due to the precise control of irrigation. Although the United States of America (USA) has a small rice production compared to Asian countries, the country is responsible for 5% of all rice exports in the world and has tripled its imports since 2001/02, showing a clear increase in crop demand (USDA, 2023).

The United States has four major rice-producing regions produced through the irrigated rice crop system: the Arkansas Grand Prairie, Mississippi Delta (Arkansas, Mississippi, Missouri, and Northeast Louisiana), Golf Coast (Texas and Southwest Louisiana), and Sacramento Valley California. These regions produce different types of rice classified by the United States, largely defined by grain market classes, with approximately 75% of the country producing long grain, 24% producing medium grain, and just 1% producing short grain. These last two types are produced partially by the state of Arkansas and mainly by the state of California (USDA, 2023).

Among all the breeding programs in the country, the Louisiana State University AgCenter is a centennial rice breeding program that utilizes a wide network of locations to conduct its METs across the US Rice belt. Because of its extensive field phenotypic evaluations, some questions were raised, such as: Is it necessary for so many locations? Does our MET match the TPE? Are we allocating our field trials properly? If we reduce the number of trials, we reduce the cost, but what happens with accuracy? In this context, several studies have indicated that the environmental covariates inclusion can enhance prediction accuracy (Montesinos-López et al., 2023; Moura-Bueno et al., 2021; Neyhart et al., 2022; Rogers & Holland, 2022). For instance, including environmental covariates for designing optimized training sets for genomic prediction can improve the response to selection per dollar invested by up to 145% compared to the model without environmental data (Gevartosky et al., 2023).

Therefore, we use historical data from the LSU Rice Breeding Program as a training set to address these questions and optimize the allocation of rice multi-environment trials (MET) in the USA Rice Belt via smart-climate prediction models based on historical weather and yield data, economic importance, artificial intelligence, and mixed model equations.

## 2. Material and Methods

### 2.1. Plant materials and trials

The experimental material consisted of 25 rice genotypes tested in two years, 2021 and 2022, and phenotyped for rice grain yield (t ha -1). There were 25 genotypes, 21 were of the long, and 4 were of the medium grain types. Louisiana State University, University of Arkansas, Horizon Ag, and Nutrien each provided five new genotypes. Also, five checks were included in the trials that were conducted at 19 different locations in the Mississippi Delta (Arkansas, Mississippi, Missouri and Louisiana) and Golf Coast (Texas and Southwest Louisiana), using both a spatial and randomized complete block design with 4 replicates per location. We define this main dataset as “LSU” for convenience.

### 2.2. Single trial analysis

We performed a two-stage analysis using linear mixed models to calculate grain yield BLUEs (Best Linear Unbiased Estimators) for each individual trial (location x year) in a similar way Jarquín et al. (2014) calculated. The first spatial model was calculated using the SpATS package in the R environment (version 4.3, https://www.r-project.org/):

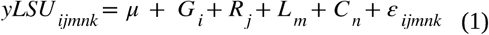

With *yLSU*_*ijmnk*_ being the phenotypic values of grain yield and equal to a overall mean *μ*, plus a fixed effect from genotypes (*G*_*i*_; *i* = 1,…,*I*), a random effect from replicates (*R*_*j*_;*j* = 1,…,*J*), a random effect from rows (*R*_*j*_*;j=l*, …, *J*), a random effect from columns (*C*_*n*_;*n*=1,…, *N*) and a random term describing residuals (ε_*ijmnk*_;*k*=1, …, *r*_*ijmn*_). The random terms were assumed to be independent and identically distributed, where 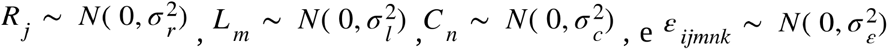. We utilized an auxiliary function in the SpATS model for spatial correction, which models the spatial or environmental effect using a two-dimensional penalized tensor-product of B-spline basis functions.

### 2.3. Multi trial analysis

The second step of the analysis was to calculate the genotype BLUPs in a multi trial analysis. This time we used *sommer* package to perform linear mixed model calculations in the R environment (Covarrubias-Pazaran, 2016):

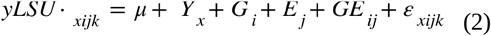

With *yLSU*_*xijk*_ being the grain yield BLUEs from model (1) and equal to a overall mean *μ*, plus a fixed effect from year (*Y*_*x*_;*i* = 1, …, *X*) as we just have two levels in this factor, a random effect from genotypes (*G*_*i*_; *i=*1, …, *X*), a random effect from environment (*L*_*j*_; j = 1, …, *J*), a random effect from the interaction between genotype and environment and a random term describing residuals *(ε*_*xijk*_; *k* = 1, …, *r*_*xij*_). The random terms were assumed to be independent and identically distributed, where 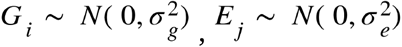 and 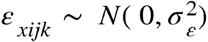. As the design matrix of the genotype by environment interaction is the Hadamard product (⊙) of [*Z*_*g*_ *Z ′*_*g*_] and [*Z*_*e*_ *Z ′*_*e*_], and *Z*_*g*_ and *Z*_*e*_ are the incidence matrix of genotype and environment,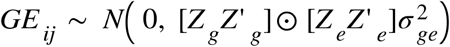.

### 2.4. Environmental covariates

The “envRtype” package was used to obtain environmental data (Costa-Neto et al., 2021). The package uses data from NASA’s orbital sensors along with location, geographic coordinates, and time range data to extract environmental data related to the experimentation locations. After obtaining the data, we tuned the environmental covariates (EC) with the cardinal limits for temperature on the phenology development of rice (Table 1). The resulting centralized and scaled matrix had 114 covariates (19 location covariates x 6 phenological stages) for 19 experimentation sites.

**Table 1.**
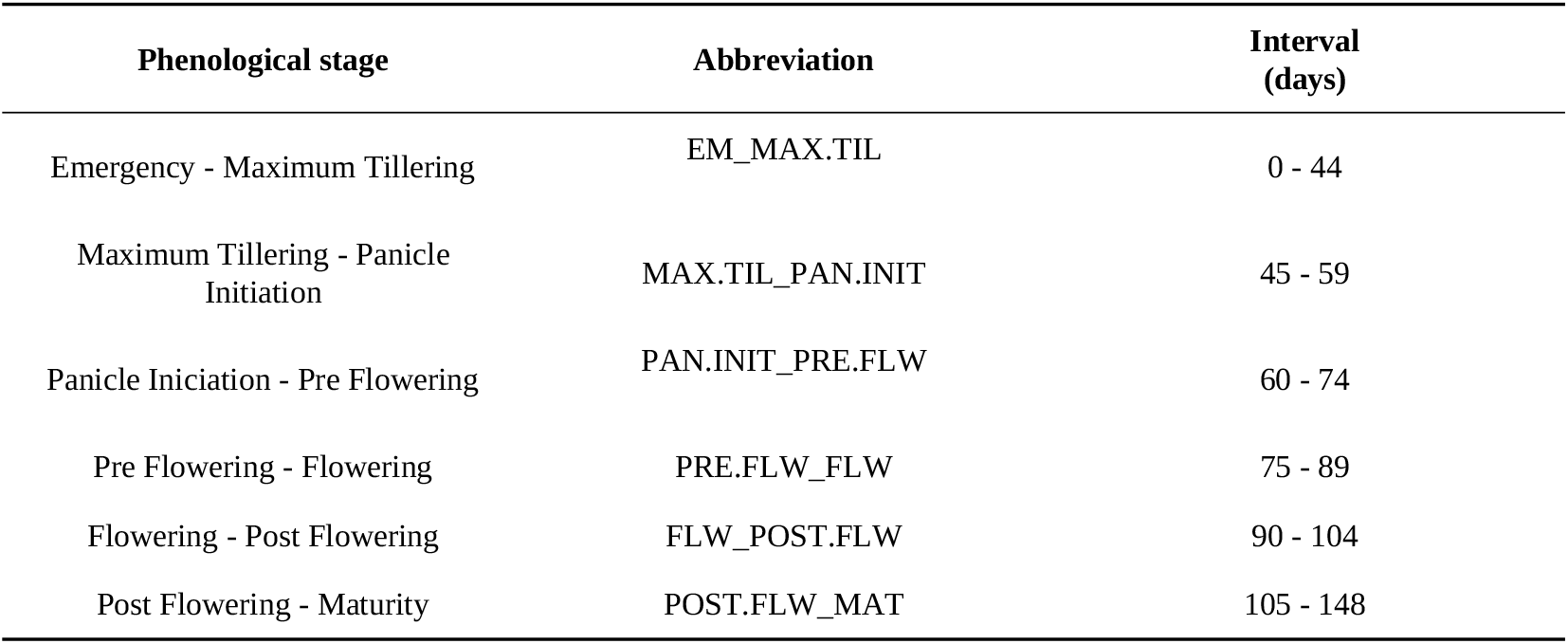
Cardinal limits for temperature on the phenology development of rice.

| Phenological stage | Abbreviation | Interval (days) |
| --- | --- | --- |
| Emergency - Maximum Tillering | EM_MAX.TIL | 0 - 44 |
| Maximum Tillering - Panicle Initiation | MAX.TIL_PAN.INIT | 45 - 59 |
| Panicle Initiation - Pre Flowering | PAN.INIT_PRE.FLW | 60 - 74 |
| Pre Flowering - Flowering | PRE.FLW_FLW | 75 - 89 |
| Flowering - Post Flowering | FLW_POST.FLW | 90 - 104 |
| Post Flowering - Maturity | POST.FLW_MAT | 105 - 148 |

The *SoilType* package was used to generate the soil covariates matrix (Fritsche-Neto, 2023). The package uses GPS coordinates to capture information about the soil of the locations closest to the experimentation site through the World Soil Database (WoSIS) (Batjes et al., 2017). Based on this database, the package also calculates several chemical and physical soil covariates. For the following steps, we joined the location and soil matrices into a single environment covariates matrix, which had 125 covariates for 18 locations, since it was not possible to generate environmental information for one of the 19 locations. This matrix underwent quality control using the *caret* package to ensure that only covariates with less than 95% correlation were maintained and to reduce collinearity between covariates (Kuhn, 2008). After this control, our scaled and centered *W* matrix remained with only 67 covariates.

### 2.5. Feature selection and clustering

We used the Recursive Feature Elimination (RFE) algorithm and the random forest learning method (Kuhn, 2008) of the caret R package to select the most important predictors or covariates. The validation method was the repeated cross-validation with 5 folds and 5 replicates. Then, models using the *W* matrix composed with only the predictors selected in the RFE, we calculated the enviromic-based kernel for similarity among environments (*Ω*), or “environmental relationship matrix” (Costa-Neto et al., 2021):

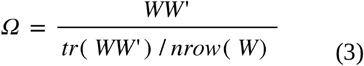

Where *W* is the environmental covariates matrix generated in the last steps, *tr* () is a trace matrix and *nrow* () is the number of rows. To group the experimental locations based on the features selected in the previous step, we clustered the locations using the *factoextra* package and k-means method (Kassambara & Mundt, 2020). Then, we decided how many clusters would be used to separate the locations with the help of the “Within Cluster of Squares” method.

### 2.6. Multi-Environment Trial Optimization

To explore the benefits of multi environment trial optimization through environmental covariates, we estimated the BLUPs for each genotype in four different scenarios:

#### I The Multi-Environment model

The Multi-Environment model (MET) is exactly the complete model (2) previously used in the MET analysis:

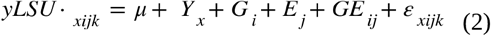

#### II The Multi-environment model with Environmental Covariates

The Multi-Environment model with Environmental Covariates (MET_EC) has the same effects as the previous model, but with the addition of the enviromic-based kernel for similarity among environments (*Ω*) based on the *W* matrix previously described as the EC matrix:

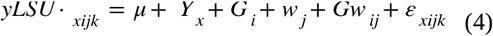

Where *w*_*j*_ is a random effect from environments (*w*_*j*_;*j* = 1,…,*J*),*Gw*_*ij*_ is a random genotype by environment interaction. With 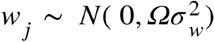 and 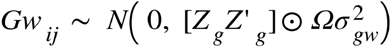.

#### III The Within Cluster Multi-Environment model

The Within Cluster Multi-Environment model (WC_MET) is the same as the MET model, but the genotype BLUPs were calculated for each of the clusters (*c*) individually (*c*=1, …, *C*) (*c*=1, …, *C*), with *C* representing the five LSU clusters with different numbers of trials each one:

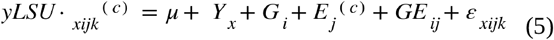

#### IV The Optimized Multi-Environment model

Finally, the Optimized Multi-Environment model (OP_MET) is the same as the MET model, but calculated individually for each subset of locations for each replicate. Where is a subset of five randomly chosen locations from each of the five clusters (*l* = 1, …, *L*) and *h* is one of the ten replicates (*h =* 1, …, *H*) performed to avoid bias by choosing only one location from each cluster:

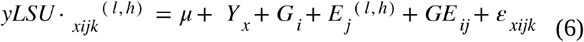

### 2.7. Cluster effect on rice grain yield

We calculated clusters adjusted means based on rice grain yield performance of each location to observe the effect of environmental clusters on productivity based on the following model:

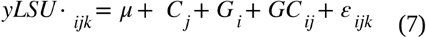

With *yLSU*·_*ijk*_being the grain yield BLUEs from model (1) and equal to a overall mean *μ*, plus a fixed effect from cluster (*C*_*j*_; *j=*1, …, *J*), a random effect from genotypes (*G*_*i*_; *i=*1, …, *I*), a random effect from the interaction between genotype and cluster (*GC*_*ij*_) and a random term describing residuals (ε_*xijk*_;*k* = 1,…, *r*_*xij*_). The random terms were assumed to be independent and identically distributed, where 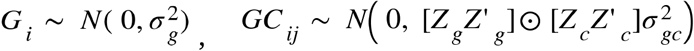, and 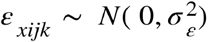, and *Z*_*g*_ and *Z*_*c*_ are the incidence matrix of genotype and cluster.

### 2.8. Accuracy per unit of dollar invested

To obtain accuracy for each of the four scenarios described above, we calculated broad-sense heritability (*H*^2^) using the Cullis method (Cullis et al., 2006):

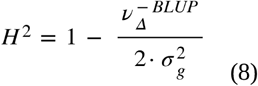

Where the term 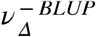 stands for the average standard error of the genotypic BLUPs. The next step was to generate the heritability value per dollar unit invested in a similar way that Gevartosky et al. (2023) calculated. For this, we assumed the actual average phenotyping costs per plot and trials, being $56,250 for the MET, MET_EC, and WC_MET scenarios and $22,500 for OPT_MET, since in the latter, the number of environments tested is almost four times smaller than the other scenarios (from 18 locations to five locations, being one from each mega-environment cluster).

### 2.9. Delimitation and economic characterization of the target population of environments

Through the previous analyses, we were able to predict the most important environmental covariates for rice grain yield by using the “LSU” MET as the training set. Based on that, we used those covariates to perform a K-means cluster analysis to delimit the target population of environments (TPE) for the whole USA Rice Belt, the South of the USA, and Louisiana. The whole USA rice belt is composed of 80 counties from seven States and represents 98% of USA rice production. Finally, based on the economic importance (rice production of each County), we estimated the mega-environment (cluster) importance and, consequently, the proportion of trials that should be allocated in that market segment. To demonstrate the environmental characterization of the regions within the US Rice Belt, we used a PCA biplot based on the W matrix, with locations separated by clusters. This plot includes a standard PCA analysis, with the addition of loading plots to show the influence of each covariate on the principal components.

We also calculated a yield-based GxE matrix (*W*_*yield*_) to understand the relationship between environments, as this is a more conventional matrix for delineating TPE and to recommend the best varieties for each TPE (Yan et al., 2023). We performed the same analyzes that we did for the *W* matrix, we calculated the yield-based environmental relationship matrix (*Ω*_*yield*_), clustered the location using the, *W*_*yield*_ calculated the trials percentage for each cluster and calculated the clusters adjusted means.

## 3. Results

### 3.1. Defining the USA rice mega-environments

The W matrix, with the ECs selected by the supervised artificial intelligence algorithm (**Figure 1A**), comprises the ones that best explain the rice yield variations across locations in the USA rice belt. The eight most critical covariates were both temperature and soil-related. Among the temperature-related variables, there was *The dew/frost point temperature at 2 meters above the surface of the earth* (T2MDEW) in the phenological stage between Flowering and Post Flowering (FLW_POST.FLW); the T2MDEW in the phenological stage between Pre Flowering and Flowering (PRE.FLW_FLW); the TM2DEW in the phenological stage between Panicle Initiation and Pre Flowering (PAN.INIT_PRE.FLW); *The minimum hourly air (dry bulb) temperature at 2 meters above the surface of the earth in the period of interest* (T2M_MIN) in the stage between Maximum Tillering and Panicle Initiation (MAX.TIL_PAN.INIT); the *Growing Degree-Days* (GDD) at the PRE.FLW_FLW stage; And *The minimum and maximum hourly air (dry bulb) temperature range at 2 meters above the earth’s surface in the period of interest* (T2M_RANGE) in MAX.TIL_PAN.INIT. Regarding the soil-related covariables, the chemical soil feature *Calcium carbonate total equivalent in g/kg* (TCEQ); and the physical soil feature *Total Silt in g/100g* (SILT). Therefore, the only phenological stages that do not produce environmental variability in rice yield are the Emergency to Maximum Tillering stage (EM_MAX.TIL), between 0 to 44 days, and Post Flowering to Maturity (POST.FLW_MAT), between 105 to 148 days.

**Figure 1.**
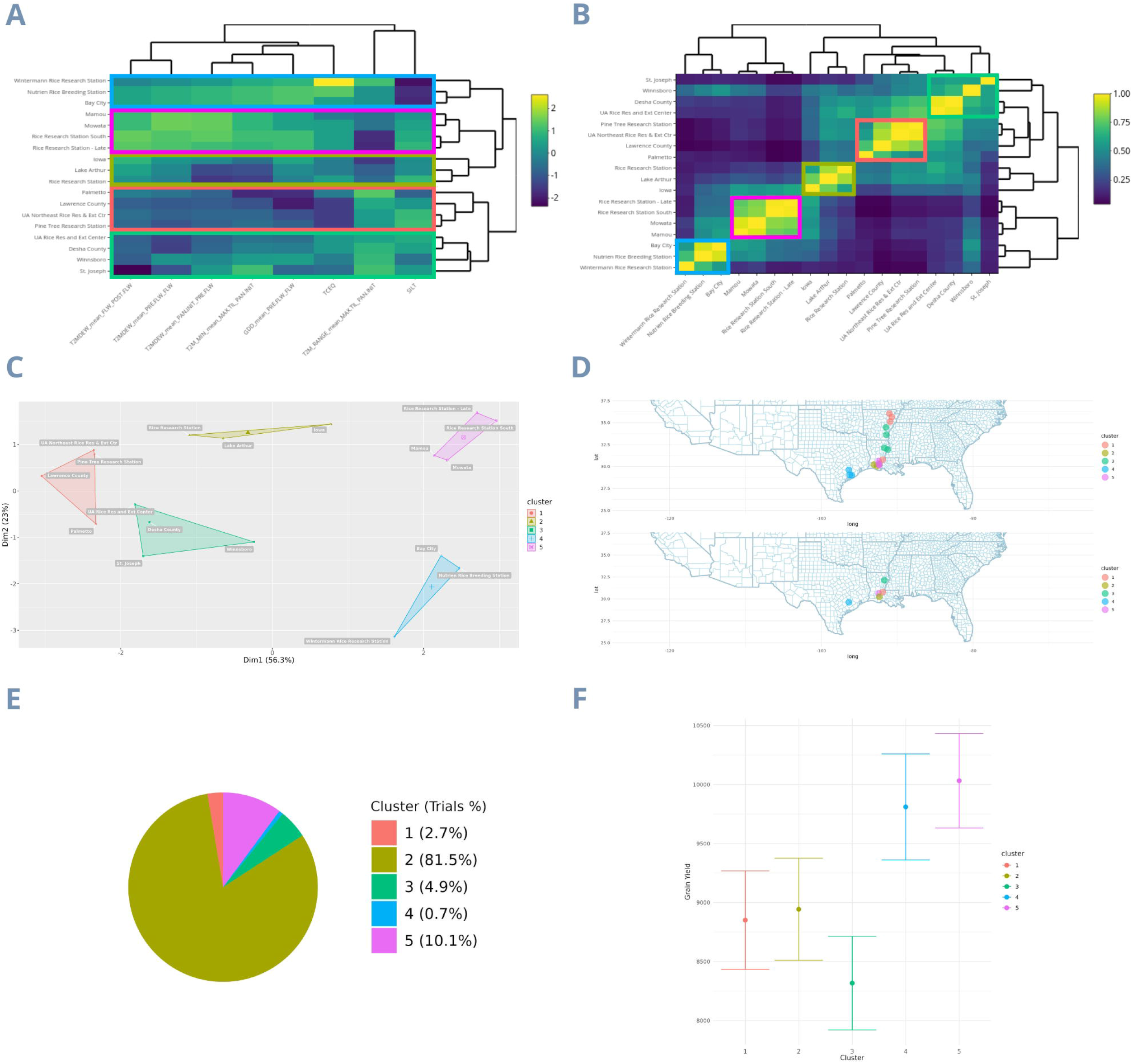
Clustering and characterization of mega-environments from the LSU dataset using the environmental covariates matrix. A - The environmental covariates matrix; B - Environmental relationship matrix; C - Clusters defining the dataset mega-environments; D - All locations on the top map and only one location per cluster after optimization on the bottom map; E - Trials percentage in each cluster; F - Cluster BLUEs. The colors of each cluster are the same in all images.

**Figure 2.**
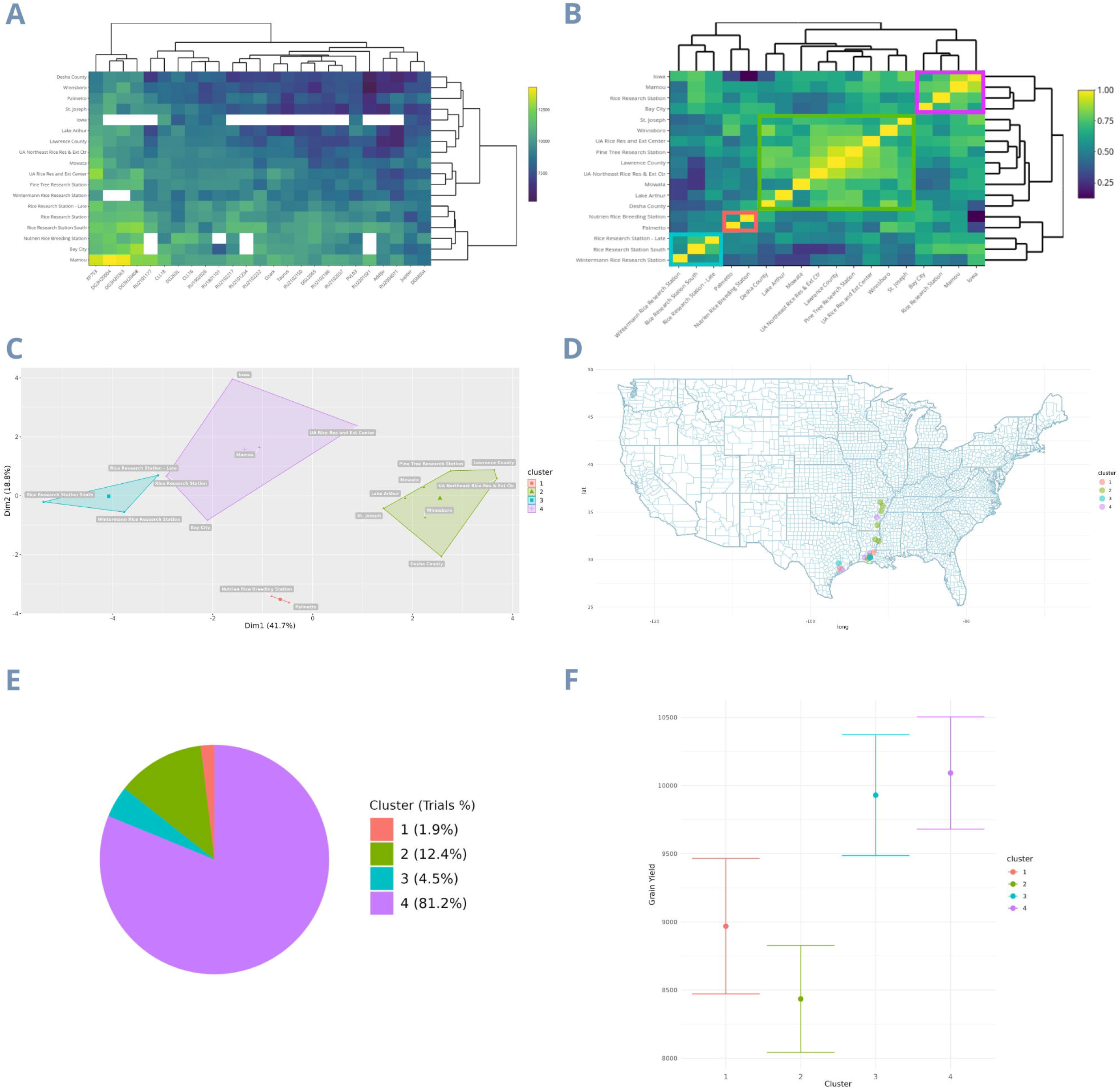
Clustering and characterization of mega-environments from the LSU dataset using the yield-based GxE matrix. A - The GxE matrix; B - Environmental correlation matrix; C - Clusters defining the dataset mega-environments; D - All locations separated by cluster on the map; E - Trials percentage in each cluster; F - Cluster BLUEs. The colors of each cluster are the same in all images.

From the *W* matrix (**Figure 1B**), we performed the clustering and thus identified the mega-environments present in the LSU rice breeding program (**Figure 1C**). This k-means clustering analysis revealed five distinct mega-environments among the eighteen experimentation sites, with the first two Principal Components (PCs) explaining 79.3% of the variation (PC1 accounting for 56.3% and PC2 for 23%). The map at the top displays the non-optimized experimentation locations, while the map optimized by the k-means method appears at the bottom (**Figure 1D**). With this optimization, the number of locations was reduced by 27.7%, covering only the states of Louisiana and Texas. Furthermore, we display the percentage of trials per cluster (**Figure 1E**), allowing us to observe the distribution of trials across mega-environments. The mega-environments represent from as little as 0.7% of total trials (Cluster 4) to as much as 81.5% of total trials (Cluster 2). Finally, we present the adjusted means for each cluster (**Figure 1F**), with Clusters 4 and 5 having the highest averages (comprising 0.7% and 10.1% of trials, respectively), with approximately 10,000 and 9,500 in Grain Yield. Included within these two clusters are the locations of Rice Research Station - Late, Rice Research Station South, Mamou, Morata, Bay City, Nutrien Rice Breeding Station (El Campo), and Wintermann Rice Research Station.

The *W*_*yield*_ matrix, which correlates with the *W* matrix by 52.59%, represents a yield gradient (**Figure 1A**). Consequently, the *W*_*yield*_ does not exhibit clearly separated kinship blocks (**Figure 1B**), and the four clusters appear closer to each other, while the variation within clusters increases. Furthermore, the first two principal components accounted for a smaller proportion of the total variation, with the first two PCs explaining 60.5% of the total variation (PC1 at 41.7% and PC2 at 18.8%) (**Figure 1C**). Regarding the percentage of trials per cluster, clusters 4 and 2 have the highest trial allocations, with 81.2% and 12.4% respectively (**Figure 1E**). The clusters with the highest yield adjusted means were 4 and 3, which include the locations Iowa, Mamou, UA Rice Research and Extension Center, Rice Research Station - Late, Bay City, Rice Research Station, Wintermann Rice Research Station, and Rice Research Station - South (**Figure 1F**).

To show the difference between the environmental relationship matrix and the yield-based GxE correlation matrix (*Ω* and *Ω*_*yield*_), we plotted a density distribution graph (**Figure 3**). The *Ω* distribution has a high density where environments have lower relationship (Peak in 0.15) and a higher standard deviation (0.28), highlighting the environmental heterogeneity contained in the MET. On the other hand, the *Ω*_*yield*_ has a high density of more correlated environments (Peak in 0.67) and a lower standard deviation (0.17), masking the real MET environmental heterogeneity.

**Figure 3.**
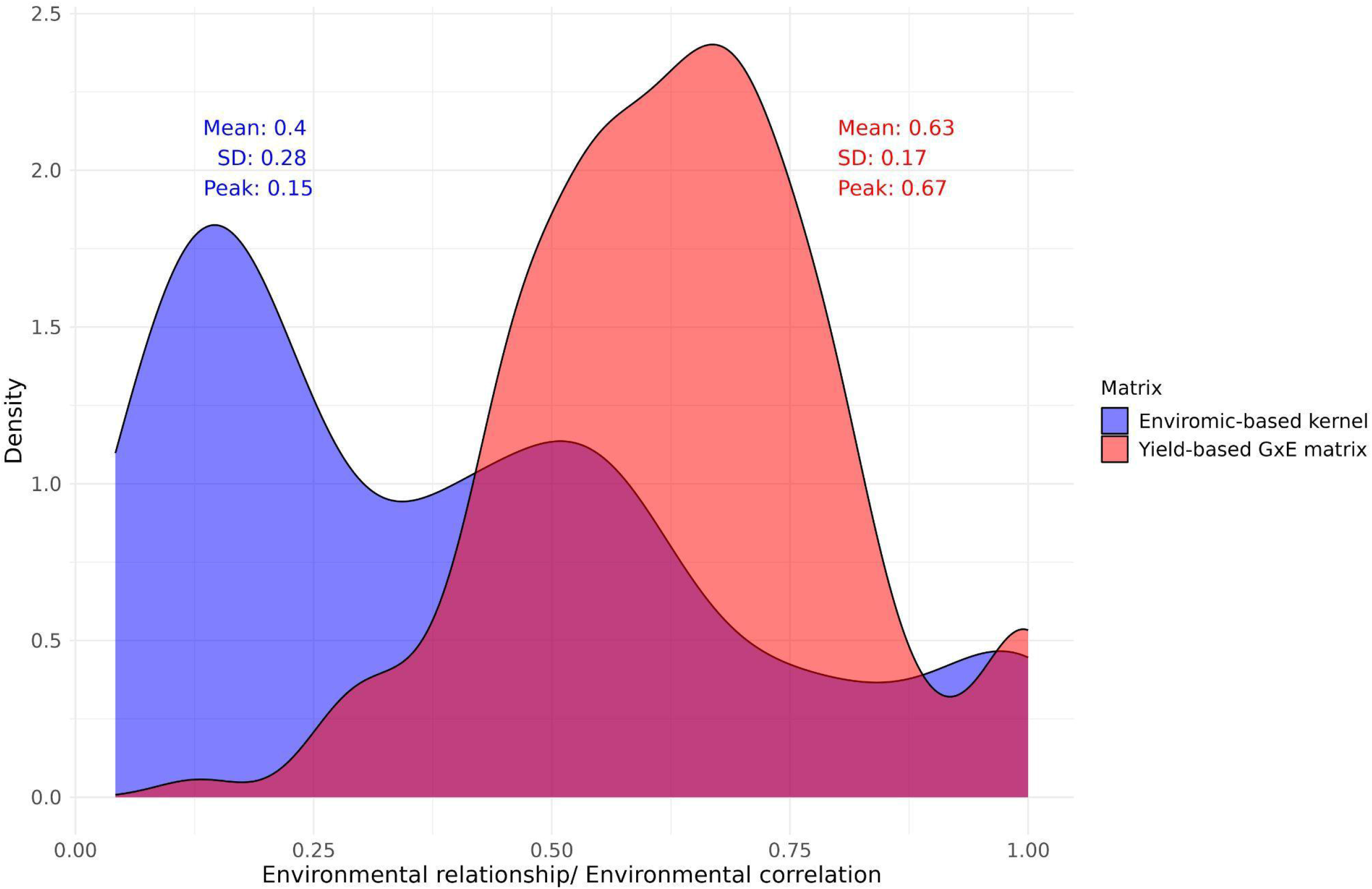
Density distribution graph of the enviromic-based kernel and yield-based GxE matrix. In blue is the distribution, mean, standard deviation (SD) and correlation value when the distribution reaches the maximum density (peak) of the enviromic-based kernel. In red are the same statistics, but for the yield-based GxE matrix.

### 3.2. Multi-Environment Trials Optimization

The heritability graph reveals distinct outcomes for the four scenarios analyzed (**Figure 4**). The MET scenario has a heritability of 0.95; the MET_EC scenario has a heritability of 0.95; the OPT_MET scenario has a mean heritability of 0.84; and the WC_MET scenario has a heritability mean of 0.75. When we divided the heritability by the total cost associated with each scenario, the ratio between heritability and cost of the OPT_MET scenario was more than twice as large as that of any other scenario.

**Figure 4.**
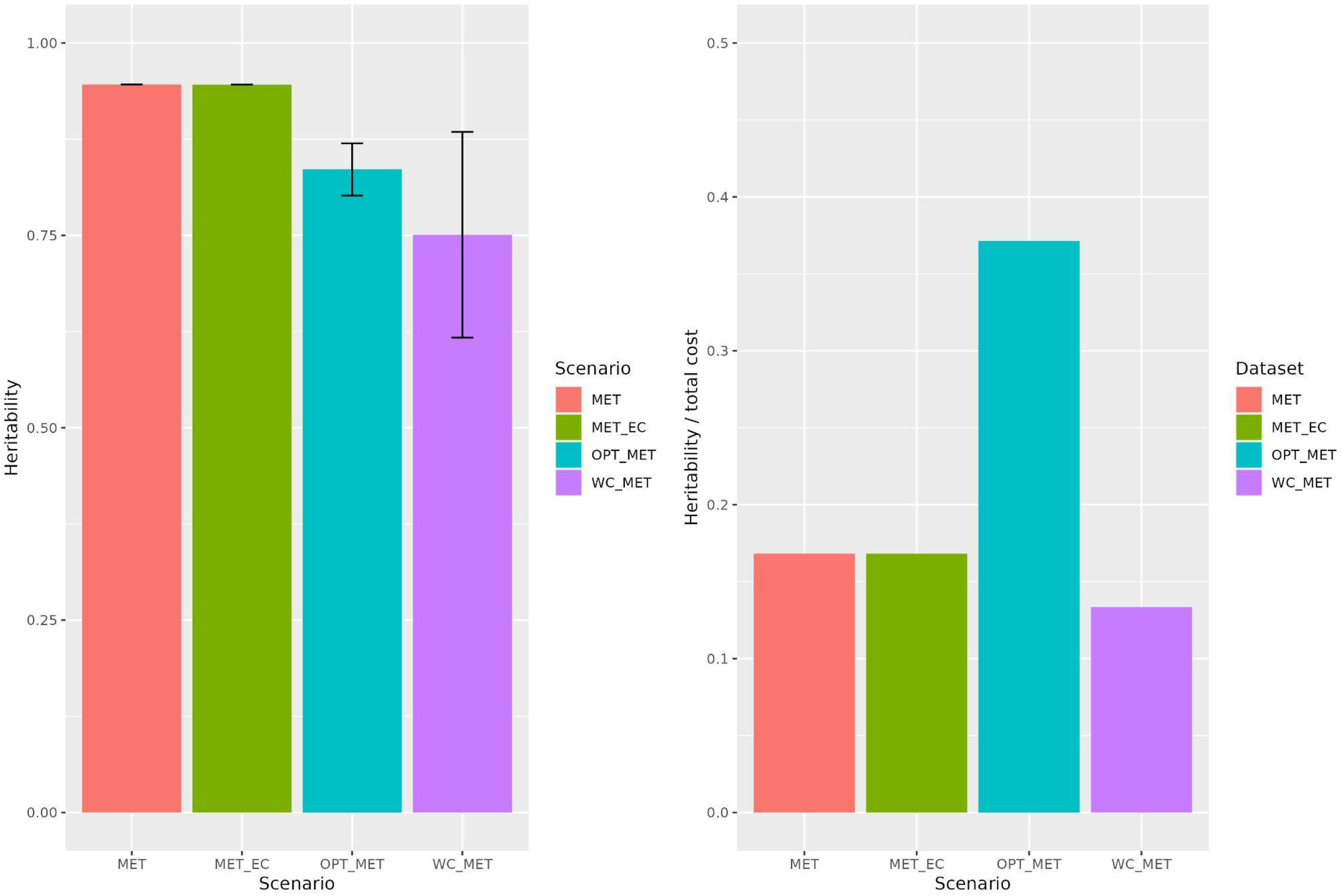
Broad-sense heritability on the left and heritability per unit of dollar invested on the right in the different scenarios for multi-environment trials optimization testing. MET - The multi-environment model; MET_EC - The multi-environment model with environmental covariates; WC_MET - The within cluster multi-environment model; OPT_MET - The optimized multi-environment model.

### 3.3. Delimitation and characterization of the USA rice belt target population of environments

The subsequent analyses delineate the mega-environments using predicted covariates that best explain rice yield from the LSU dataset (**Figure 5**). We display the clustering and distribution of trials across all 80 counties, representing 98% of US rice production in Arkansas, California, Florida, Louisiana, Mississippi, Missouri, and Texas (**Figure 5**). The *Ω* matrix highlights a distinct group unrelated to the other counties and comprises those in California (**Figure 5B**). Consequently, the clustering of the U.S. dataset revealed significant variability between the California cluster and the rest of the American counties, resulting in the latter being considered a single cluster. This created a new dataset that excluded the 9 California counties and utilized 85.6% of the remaining trials (South dataset). (**Figure 5E**). The findings from this dataset indicated that the revised W, which includes the 71 remaining counties, is more homogeneous and is divided into four mega-environments (**Figure 6C**). The distribution of trials across these clusters ranges from 12.7% to 44.4% (**Figure 6E**). Lastly, a new dataset comprising 19 Louisiana parishes was divided into two mega-environments (**Figure 7C**), representing Louisiana’s northern and southern regions (**Figure 7D**), with 27.7% and 72.3% of trials per cluster, respectively.

**Figure 5.**
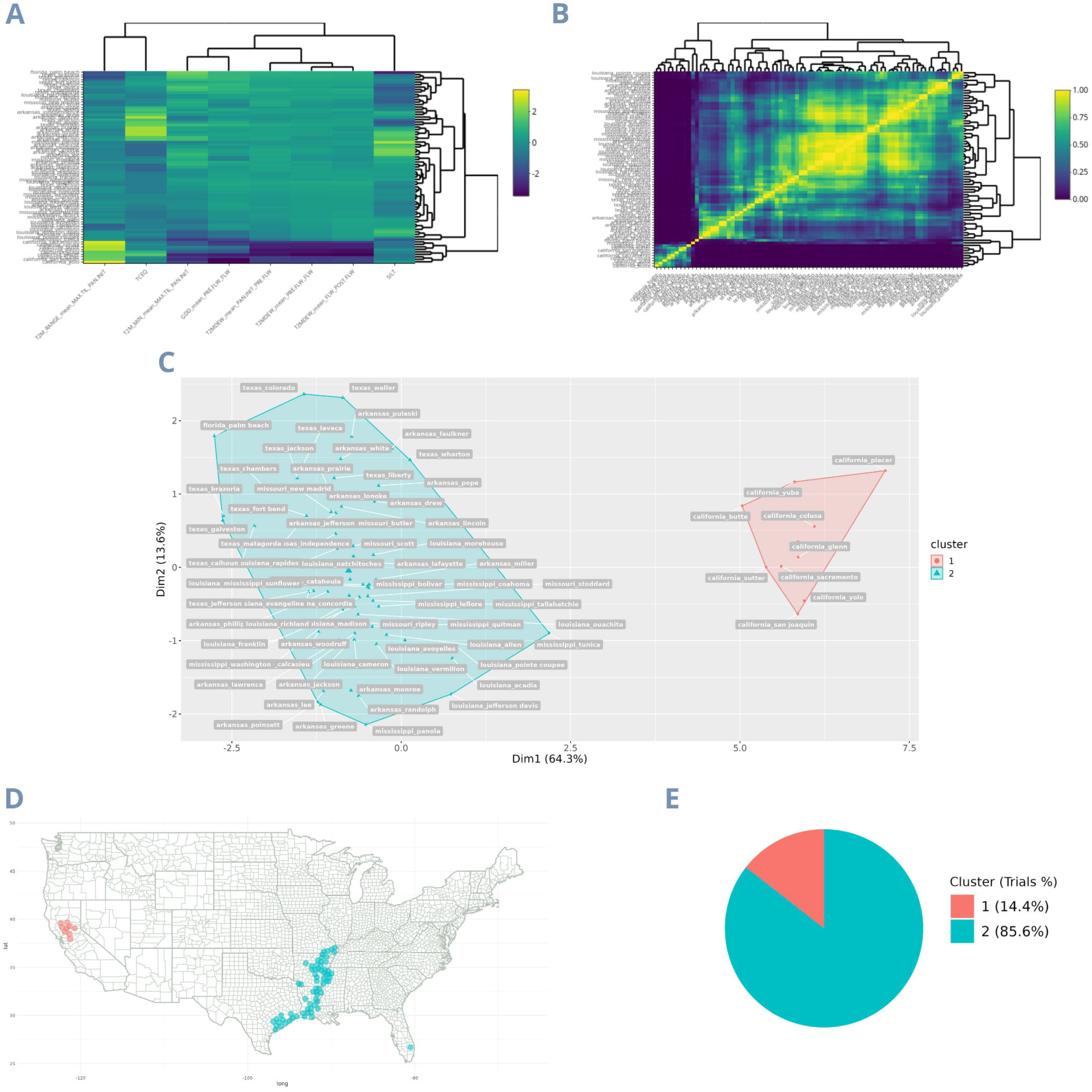
Clustering and characterization of mega-environments from the USA dataset using the environmental covariates matrix. A - The environmental covariates matrix; B - Environmental relationship matrix; C - Clusters defining the dataset mega-environments; D - All locations separated by cluster on the map; E - Trials percentage in each cluster. The colors of each cluster are the same in all images.

**Figure 6.**
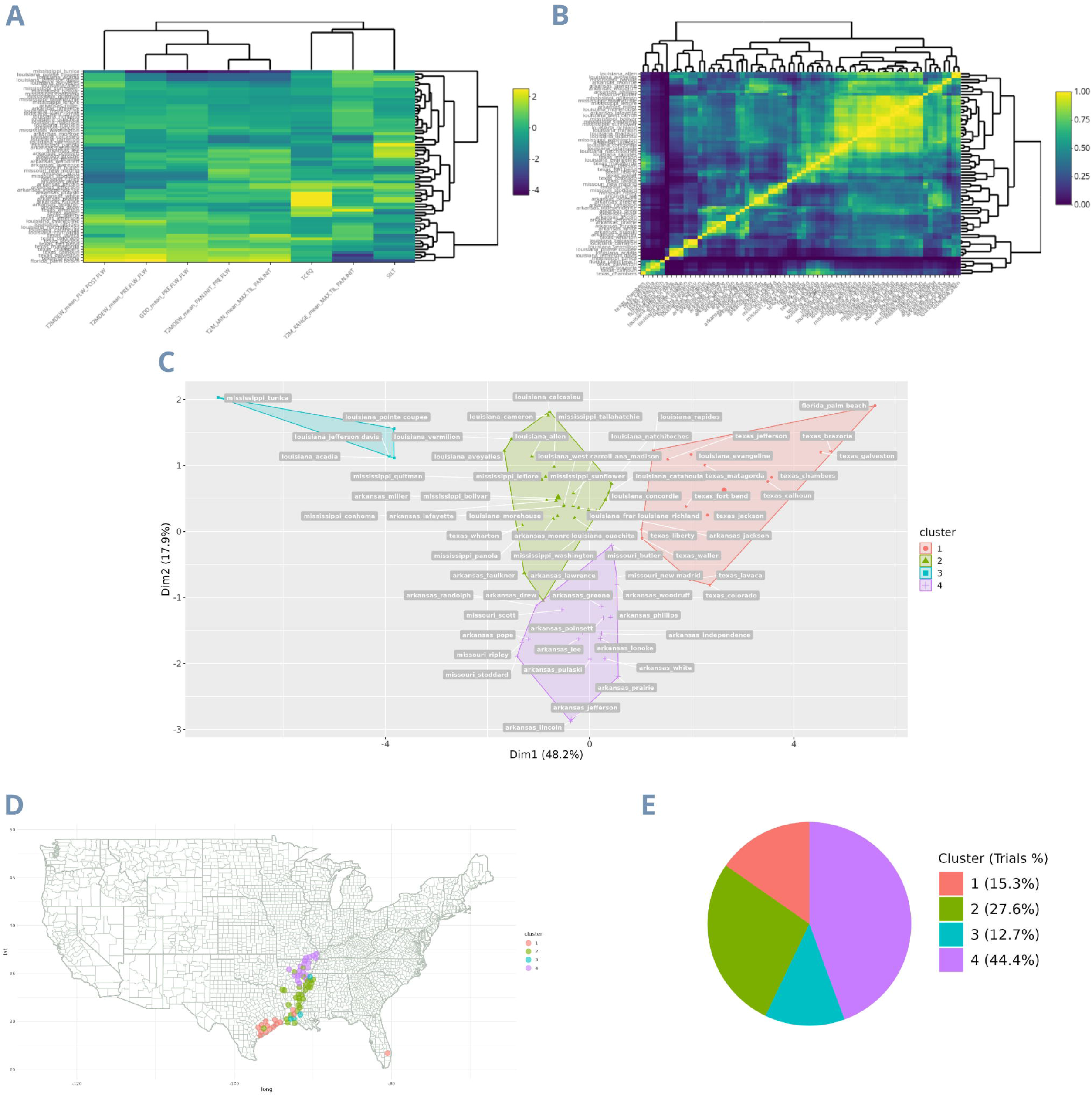
Clustering and characterization of mega-environments from the South dataset using the environmental covariates matrix. A - The environmental covariates matrix; B - Environmental relationship matrix; C - Clusters defining the dataset mega-environments; D - All locations separated by cluster on the map; E - Trials percentage in each cluster. The colors of each cluster are the same in all images.

**Figure 7.**
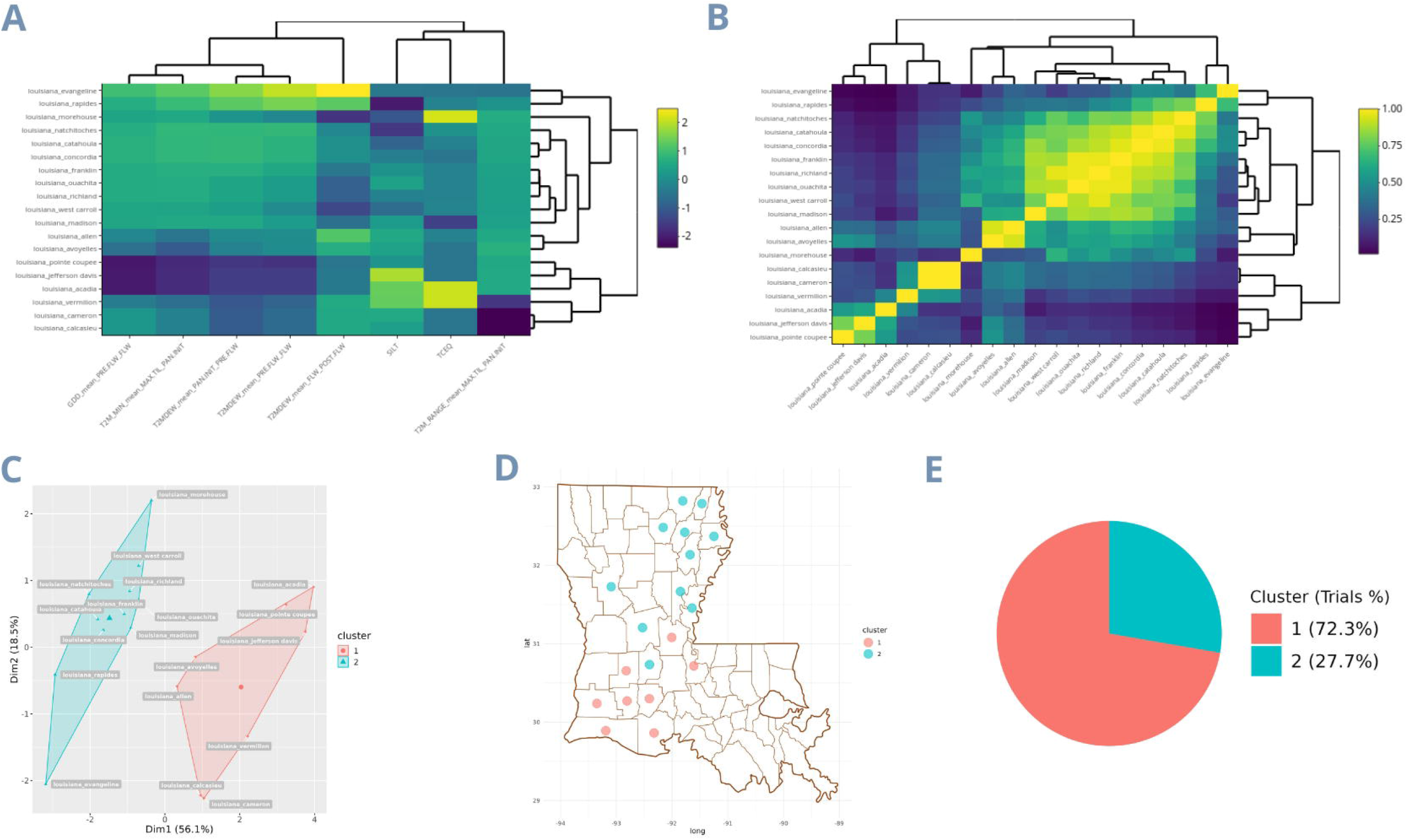
Clustering and characterization of mega-environments from the LA dataset using the environmental covariates matrix. A - The environmental covariates matrix; B – Environmental relationship matrix; C - Clusters defining the dataset mega-environments; D - All locations separated by cluster on the map; E - Trials percentage in each cluster. The colors of each cluster are the same in all images.

LSU’s environments of evaluation were divided into five clusters, and these clusters are characterized based on the evaluation of PCA biplot analysis (**Figure 8**):

**Figure 8.**
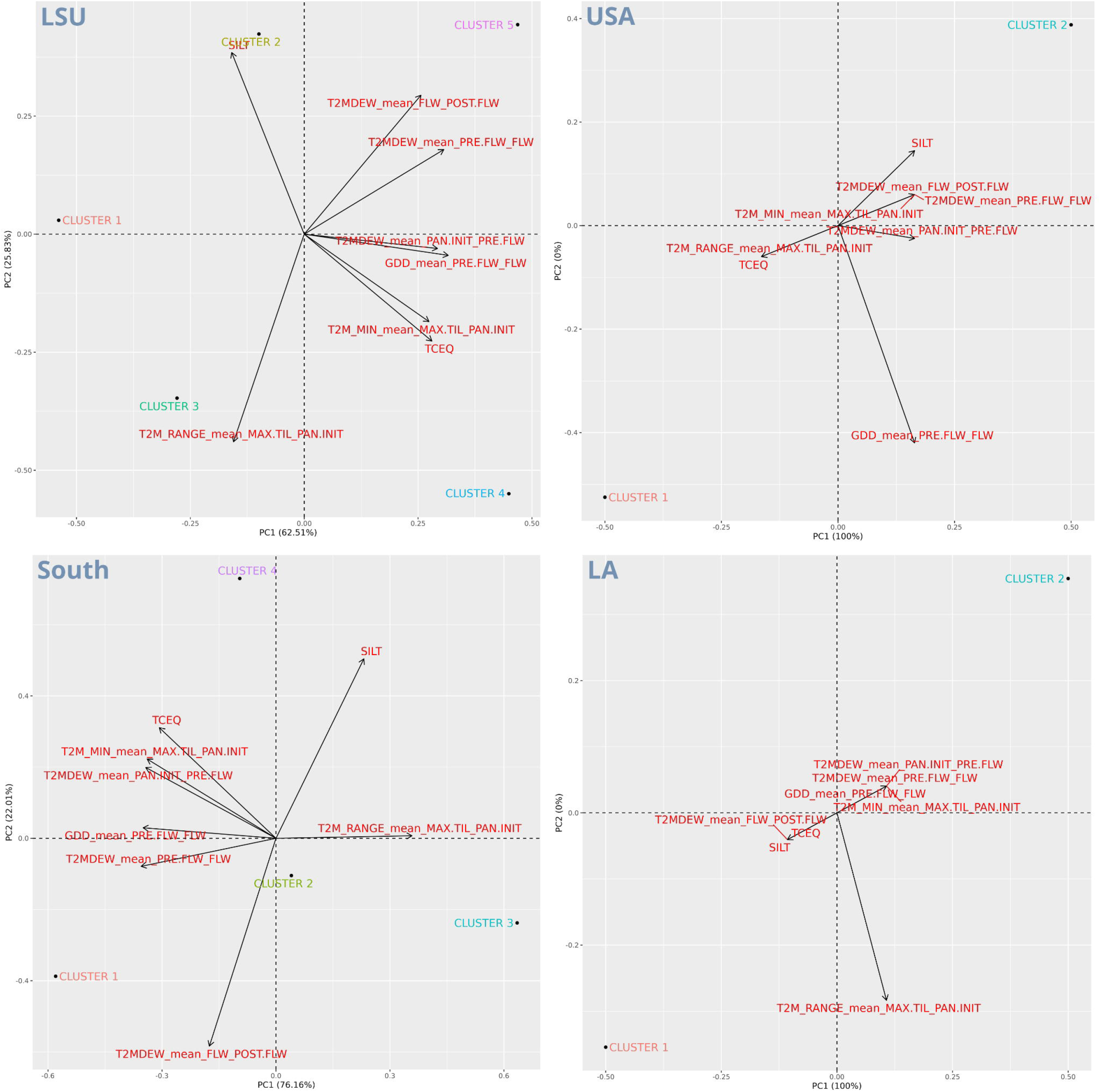
Characterization of rice yield United States target population of environments. For each of the four PCAs biplots, the separate dots represent the clusters, the black dashed lines divide the four areas of the graph with respect to the two main components that most explain the variance of the clusters, the size and direction of the black arrows show how much the variables (in red) contribute to the variance of the main components. The closer the clusters (black dots) are to the variables, the more related the clusters are to high values of that variable.

- Cluster 1 is related to lower covariates values, such as minimum temperatures (MAX.TIL_PAN.INIT), dew point temperatures (from PAN.INIT to POST.FLW), and TCEQ.
- Cluster 2 exhibits higher silt contents and lower minimum temperatures (MAX.TIL_PAN.INIT), dew point temperatures (T2MDEW), and TCEQ.
- Cluster 3 is characterized by higher temperature ranges (MAX.TIL_PAN.INIT), while lower values for all the other covariates.
- Cluster 4 has higher minimum temperatures (MAX.TIL_PAN.INIT) and TCEQ, lower dew/frost temperatures (PRE.FLW to POST.FLW), and SILT content.
- Cluster 5 exhibits higher TCEQ content and minimum temperature (MAX.TIL_PAN.INIT), while SILT and dew/frost point temperatures have lower values.

## 4. Discussion

MET are necessary to study the phenotypic plasticity of the genotypes when subjected to environmental fluctuations under different crops’ phenological stages. The change in the genotypes rank can result from more static and predictable environmental covariates, such as SILT and TCEQ as observed in our study, but also from more unpredictable ones, such as temperature in our study (Crespo-Herrera et al., 2021). The evaluation of quantitative traits that are highly prone to demonstrate phenotypic plasticity (crossover GxE interactions) must be done so that selection sites effectively meet TPE needs. The assumption is that the locations chosen to allocate the trials will efficiently represent the sets of environmental covariates found in the TPE to make it possible to outline a strategy on how to handle GxE in the selection of superior genotypes in different stages of a breeding program (Cooper & Delacy, 1994). In this context, disregarding the GxE interaction can lead to a reduced selection response. This is particularly true when programs conduct early generation selection, as they test many lines in few environments (Cooper et al., 1995). Therefore, information on the characterization and design of TPEs can benefit breeding programs by guiding the allocation of trials and considering the alignment of selection environments and TPEs.

Our study found eight covariates that explain 58% of all the rice yield variation in the USA Rice Belt. The environmental covariate that explains the largest variation is temperature. Although it is widely known that rice flowering is regulated by temperature, little is still known about its regulation mechanism. Generally, higher minimum temperatures accelerate crop flowering, reducing biomass and grain yield. However, late flowering may induce biomass accumulation and concomitantly reduced grain filling (Srikanth & Schmid, 2011). Also, high temperatures can deteriorate rice quality due to an imbalance between protein content and starch in the grains (Liu et al., 2021). On the other hand, cold stress can also reduce rice yield during any phenological stages (Cruz et al., 2013). In our study, the minimum temperature was particularly important between maximum tillering and panicle initiation, with higher minimum temperatures representing clusters with higher yields. Moreover, some studies highlight the importance of assessing air relative humidity when studying the response of rice to temperature (Stuerz & Asch, 2019). In our study, the dew/frost temperature, which depends on air humidity and temperature, was a key covariate to explain the different selection environments. Besides the covariates mentioned above, two soil covariates were also responsible for the yield change in the trials, the amount of calcium and silt.

Using these covariates, we grouped the locations into five clusters with environmental similarities. The productivity of these groups ranged from 8500 to 10000 tons per hectare for clusters 3, 1, 2, 4, and 5, respectively. The two clusters with the highest average productivity were associated with high minimum temperatures, dew point temperatures, and high TCEQ contents in the soil. However, the two clusters with the greatest economic importance in the American TPEs are clusters 4 and 2 from the South dataset, comprising 44.4% and 27.6% of the entire rice production in the USA Rice belt (**Figure 6D** and **Figure 6E**). These same clusters are located in southern Missouri, Arkansas, and northern Louisiana (in green and purple). In the LSU MET map, these locations correspond to clusters 1 and 3 (in red and green), with only 2.7% and 4.9% of the trials in that region. The results of the delimitation and characterization of TPEs demonstrate how trial allocation could be optimized for more efficient resource utilization. Additionally, the results highlight the potential for improvement in US rice TPE varieties, as environments with greater economic importance (Cluster 1 and 3 in LSU) have lower productivity and less favorable environmental covariates (**Figure 8**), such as lower minimum temperatures and dew/frost points. Furthermore, when covariates are deemed highly significant for a trait to the extent that they become a long-term breeding target, this would justify the establishment of fine-grained research facilities for breeding programs (Cooper & Messina, 2021; Crespo-Herrera et al., 2021). There’s the possibility to conduct coarse-grained phenotyping, such as environments with and without a specific stress, or fine-grained phenotyping, encompassing a broad range of this environmental continuum and allowing for detailed study of genotype reaction-norm models (Cooper & Messina, 2021).

Furthermore, our study shows, in practice, other ways to use enviromics to perform a good allocation of resources. We tested both the heritability increase with the addition of covariates in MET joint analysis and the optimization of MET by using a reduced and efficient number of sites. Although the OPT_MET scenario did not produce the highest heritability among all scenarios, it was possible to maintain a high level per unit of dollar invested. Indeed, this value was more than twice as large as any other scenario (**Figure 4**). Testing numerous lines in many environments for years can lead to a good genotype performance recommendation. However, this strategy assumes that the same genotypes will be used in the future and that there is no budget limit, which is inaccurate for a breeding program. In this sense, we reduced the number of tested locations by 27.7% while maintaining high accuracy (0.8357). Reducing the number of tested lines and allocating them according to the economic importance of each mega-environment will lead to lower costs for breeding, mainly in one of the costliest stages in the process, the phenotyping.

In recent years, breeders have shown promising results when considering environmental covariates in data modeling. Gevartosky et al. (2023) improved the response to selection by 145% when they considered environmental covariates in the design of optimized training sets for genomic prediction. In the same way, Costa-Neto et al. (2023) achieved a GxE variance decrease from 22% to 15% when environmental covariates were considered, showing a more effective GxE effect capture. Conversely, the heritability of the MET and MET_EC scenarios (0.9 5 and 0.9 5, respectively) remained the same even when using the *Ω* matrix in the last scenario. We believe this may have happened between the two scenarios due to the similarity between the *E*_*j*_ and *w*_*j*_ effects (where 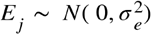, and 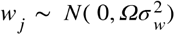). Probably, the 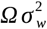 matrix (MET_EC) didn’t provide any additional information to the model to boost accuracy, as the 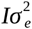 matrix (MET) already had enough information to produce the highest heritability of all scenarios. This may have occurred because the variation between locations within clusters was very small compared to the variation between clusters (**Figure 1C**).

Spindel & McCouch (2016) showed that the more correlated the environments and genotypes are in the modeling, the higher the accuracy. Therefore, it was expected that the WC_MET scenario would perform similarly to the other scenarios, but its average was the lowest (0.75 *±* 0 .1336). However, WC_MET also had the highest standard deviation, with the lowest heritability being from cluster 2 (0.54), which is the biggest (81.5% of trials). The large number of trials in this cluster may have led to reduced accuracy compared to other clusters, since the cluster locations appear highly correlated in the *Ω* matrix (**Figure 1B**).

Besides proving that environmental covariates can help better trial allocation while maintaining accuracy, we wanted to highlight the benefits of delineating mega-environments by comparing *W* and *W*_*yield*_ matrices. The traditional way to represent GxE and the correlation between environments is through the use of the *W*_*yield*_ matrix (Cooper & Delacy, 1994; Senguttuvel et al., 2021; Yan et al., 2023), as the Additive Main Effects and Multiplicative Interaction (AMMI) model (Gauch Jr. & Zobel, 1997). Using this methodology, it is possible to discover which environment produces the highest yield and which variety is better for each environment, as long as they are tested in those same environments. However, when the *W* matrix is used, it is possible to know which genotypes perform well for each environmental covariate or set of covariates due to environmental stratification. Just having information from an entire environment means the breeder cannot expand this information to new environments. The advantage of this is that environmental information is broad and free (Costa-Neto et al., 2021), while the traditional GxE matrix is highly dependent on the genotypes and environment combinations used. Therefore, with historical weather data and genotypes reaction norms, it is possible to recommend the best and most stable varieties for each target region and even work to discover potential new producing regions (Araújo et al., 2024; Callister et al., 2024; Cooper & Messina, 2021; Costa-Neto et al., 2023).

As climate varies significantly from year to year, a collection of environments cannot be deemed a TPE based solely on one or a few years of data; delimitation must be conducted based on repeatable GxE patterns (Singh et al., 2006). This fact poses further challenges to establishing TPEs using yield-based environment relationship matrices. In the case of *W* matrix map (**Figure 1D)**, the clusters are regionally separated. In contrast, for the *W*_*yield*_ matrix the clusters are mixed throughout the map (**Figure 2D**), suggesting an effect not accounted for in the last matrix and consequently, a possible change of genotype ranks between environments. The fact that the *W*_*yield*_ matrix PCs explain less variation than the *W* matrix PCs also shows more variation than was captured by the first methodology. It is possible to predict this by the *W* _*yield*_ matrix structure, a yield gradient with more homogeneous correlation between environments (**Figure 3A**), and less environmental structure captured in the correlation matrix (**Figure 3B**). Consequently, the clusters are closer to each other in the *W*_*yield*_ matrix (**Figure 2C)**, while the cluster variation is bigger.

Conclusively, our study enabled us to virtually reduce 28% of the trials while maintaining almost the same accuracy. Furthermore, we demonstrated how this trial reallocation will allow for better utilization of our resources, as we could better represent all TPEs within the USA Rice belt according to their economic importance. Additionally, we identified which environmental covariates have the greatest impact on rice productivity in the considered TPEs and that they explain 58% of all variation in rice yield in the USA. With this information, it is possible to establish fine-grained phenotyping and expand production to potential new areas. These findings can be invaluable information in assisting rice breeding efforts in the USA and aiding breeders in optimizing trial allocation.

## Author Contributions

MP and RFN elaborated on the hypothesis, conducted the analyses, interpreted the results, and contributed to the writing. AF and KG elaborated on the hypothesis, interpreted the results, and contributed to the writing. All authors read and approved the final manuscript.

## Data Availability Statement/ Supplementary material

The code and data used for the analyses described in this manuscript is publicly available on GitHub: [GitHub - MET_Optimization] (https://github.com/MelinaPrado/MET_Optimization.git). This repository includes the project structure, scripts, and data necessary to reproduce the presented results.

